# Isolation of and Characterization of Neutralizing Antibodies to Covid-19 from a Large Human Naïve scFv Phage Display Library

**DOI:** 10.1101/2020.05.19.104281

**Authors:** Andy Q. Yuan, Likun Zhao, Lili Bai, Qingwu Meng, Zhenguo Wen, Yanhu Li, Daqing Guo, Shanshan Zhen, Xiaojun Chen, Ji Yang, Xiaoying Xue

**Author notes:** Correspondence: Andy Yuan, PhD., Excyte LLC, 15601 Crabbs Branch Way, W123, Rockville MD 20855 USA.

## Abstract

SARS-CoV-2 (Covid-19) has caused currently ongoing global plague and imposed great challenges to health managing systems all over the world, with millions of infections and hundreds of thousands of deaths. In addition to racing to develop vaccines, neutralizing antibodies (nAbs) to this virus have been extensively sought and are expected to provide another prevention and therapy tool against this frantic pandemic. To offer fast isolation and shortened early development, a large human naïve phage display antibody library, was built and used to screen specific nAbs to the receptor-binding domain, RBD, the key for Covid-19 virus entry through a human receptor, ACE2. The obtained RBD-specific antibodies were characterized by epitope mapping, FACS and neutralization assay. Some of the antibodies demonstrated spike-neutralizing property and ACE2-competitiveness. Our work proved that RBD-specific neutralizing binders from human naïve antibody phage display library are promising candidates to for further Covid-19 therapeutics development.

## Introduction

Around December of 2019, pneumonia patients afflicted by an unknown type of coronavirus, which later named as 2019-nCoV, SARS-CoV-2 or Covid-19 by WHO, had emerged in Wuhan China. In the past 5 months, the highly contagious virus has since then quickly spread to almost every nation in the world, infecting more than 5 million people and resulting in more than 320,000 deaths^1^. To this date, this pandemic marked the third highly pathogenic coronavirus human infection in the 21st century, after the outbreak of the deadlier Middle East Respiratory Syndrome coronavirus (MERS-CoV) and the Severe Acute Respiratory Syndrome coronavirus (SARS-CoV-1). The whole world was caught off guard to this challenge. To flatten the incident curve, social measures such as quarantine, lockdown and distancing were widely applied, leading to a sudden pause of most economic activities. Among all the measures mobilized to combat the plague, like to many others, scientific approach is the decisive and ultimate answer.

In past months biological studies have found strong conservatives and resemblances between SARS-CoV-1 and Covid-19 in genomic sequences, mode of infection and to certain extent, clinical pathology^2,3,4^. Like dealing many infectious diseases, the first effective step whenever possible is always preventing the initiation of an infection by isolation or prophylaxis measures such as antibody or vaccine. Vaccines potentiating host immune responses and eliciting protective antibody production will quickly mobilize humoral and cellular immunity defenses against subsequent viral invasion. When lack of vaccine, the passive immunity conferred by specific monoclonal antibodies has well been recognized in the treatment of many viral diseases, for example, FDA approved anti-RSV drug Synagis^5,6^. Based on the limited scientific data, treating Covid-19 patients with convalescent plasma from recovered donors appears safe, clinically effective and reduces mortality^7^. Significant efforts have been made to develop therapeutic anti-viral antibodies against SARS-CoV^8^ and MERS^9^. High viral biological similarities and promising animal efficacy of these antibodies highlight the potential to obtain Covid-19 specific neutralizing antibodies (nAbs) for its therapy and prophylaxis.

The entry of Covid-19 to human cells has been recently reported that Covid-19 virus binds to the human angiotensin-converting enzyme 2 (ACE2) through it spike protein, or S-protein, which is almost identical in the infection of SARS-CoV-1 or MERS^4, 10^. The portion in S-protein, called receptor binding domain (RBD), is the exact motif that interacts with ACE2^11^. Details of the molecular interaction between RBD and ACE2 have been analyzed to atom level^12^, facilitating the design of blockade agents, including antibody to intervene the entry process. In the meantime, in a race to win this unprecedented battle, every second matters in the generation of lead antibody molecules. Instead of immunizing animals and screening subsequent hybridoma, we took advantages of an in-house very large naïve human antibody phage display library built in our lab. Here we summarized the construction of the library, the isolation and characterization of RBD nAbs, providing strong basis for further development of potential antibody arsenals for COVID-19.

## Results

### Human antibody phage display library

Human antibody phage display library has become the dominant route^13, 14^ to quickly obtain antibody leads and develop antibody-based biotherapeutics to an almost given target. A large human naïve antibody phage display library (Named as EHL Library) was built in-house based upon conventional phage display technique, in a similar protocol reported earlier^15^. To ensure the library to be a high-value research resource, we adopted multiple measures to achieve extensive diversity and super-size during the whole process. PBMCs of 37 donors from a variety of ethnic groups and nationalities across the world, including Latinos, African American, Caucasian, Chinese, Russians, Hindus, Vietnamese, Philippines, Native American, Japanese and Jewish etc. were collected to enhance the diversity range of antibody variable genes to be used in the scFv assembly. 7 subfamilies of variable heavy (VH) genes, 7 subfamilies of variable kappa genes (Vκ) and 11 subfamilies of variable lambda genes ( Vλ) were separately amplified and recovered. Two types of scFv libraries, VH-Vκ and VH-Vλ, were separately assembled and cloned into phagemid pADL-10b, resulting two types of scFv libraries, each containing more than 5×10^10^ colony forming unit (cfu) in size, after total 100 repeats of electroporation of ligated phagemid constructs. Random sequencing of 100 colonies of each library revealed over 80% of correct scFv-coding and full usage of all antibody subfamily genes (data not shown). The utility and quality of the EHL Library were validated by successful screening of specific hits over a dozen of selected human targets, mostly tumor-associated antigens, with average KD of two digits nM (data not shown).

### Validation of Covid-19 spike-expressing cell line

A surrogate tool is essential to research on any very deadly contagious pathogens such as Covid-19, whose spike protein is of great needs not only as purified form but also as membrane one. A cell membrane protruding spike may not behave the same as it presents in its viral particle, the availability of such cell line can serve most needs to some extent such as antibody binding examination and competition assays. A mammalian cell line ID8 was generated by transfection of 293FT with a pseudo lenti vector that contains gene encoding the full-length Covid-19 spike protein, a transmembrane motif (TM) and a 3xFLAG tag at C-terminal (Fig. 1, A) and selected under Blasticidin S hydrochloride. To verify the spike expression, structure and function, ID8 cells was stained by anti-FLAG-FITC (Fig. 1, B) or human ACE2-mFc, followed by anti-mouse-FITC (Fig. 1, C). and analyzed by FACS. Results showed that both stains exhibited significant right shift of ID8 population (Fig.1, B and C, red peaks), validated the ID8’s ability to bind ACE2, indicating the spike is in trimeric membrane status and mimics the function of Covid-19 spike^16^.

**Fig. 1.**
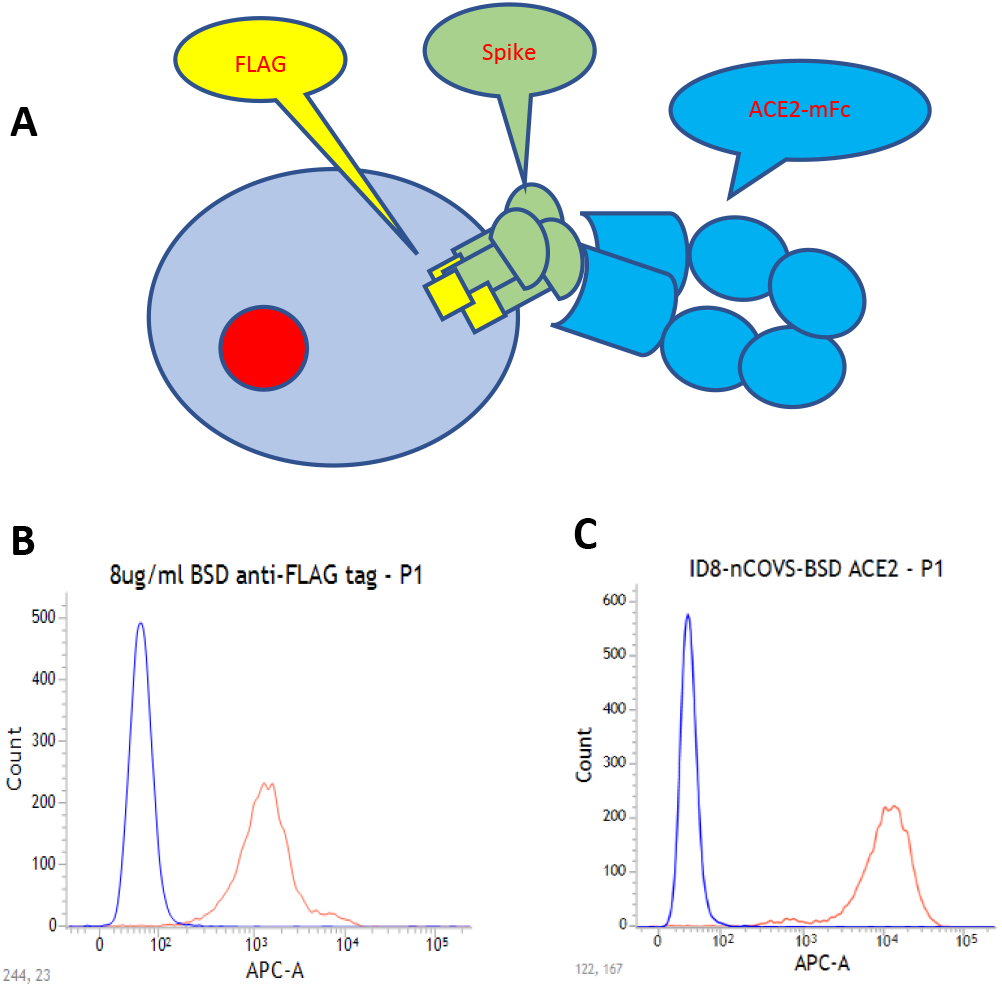
Validation of Covid-19 spike expression cell line. A mammalian cell line ID8 was generated by transfecting 293FT with a pseudo virus that contains full-length Covid-19 S protein, a transmembrane motif (TM) and a C-terminal 3xFLAG tag. **A**. The cartoon illustration of the ID8 cells and membrane anchoring of Covid-19 spike (green), C-terminal FLAG (yellow) and binding of ACE2-mFc (blue) to S trimer. **B**. FACS plot of intracellular staining of FLAG with anti-FLAG-FITC. **C**. FACS plot of ID8 cell membrane sequential staining of ACE2-mFc, anti-mouse FITC.

### Biopanning naïve phage antibody library yielded multiple RBD hits

Aiming to obtain antibodies that directly target to RBD of Covid-19 spike so that they have limited binding sites, and great chance of interference to the interaction between spike and ACE2, we used RBD fusion (RBD-mFc) instead of full-length or subunit of Covid-19 spike protein as bait in the biopanning from EHL library in a solid phase approach^17^. 3 rounds of biopanning yielded vast enrichment as output titer increased over 100 times (data not show) over that of 1^st^ round. Upon monophage ELISA screening on the 3^rd^-round output, more than 95% of clones were positive. After sequencing the scFv genes of positive hits, we found very focused clone specificity (2 uniqueness among 90 available scFv sequences, data not shown). To harvest diversified binders, we turned to 2^nd^ round output for ELISA screening (Fig2, A). Among 160 positive hits we identified 42 unique clones, with many have minor differences in amino acids. Through multi-alignment of the scFv primary sequences, a phylogenetic tree was generated (Fig 2, B) to measure homology gap between them.

**Fig. 2.**
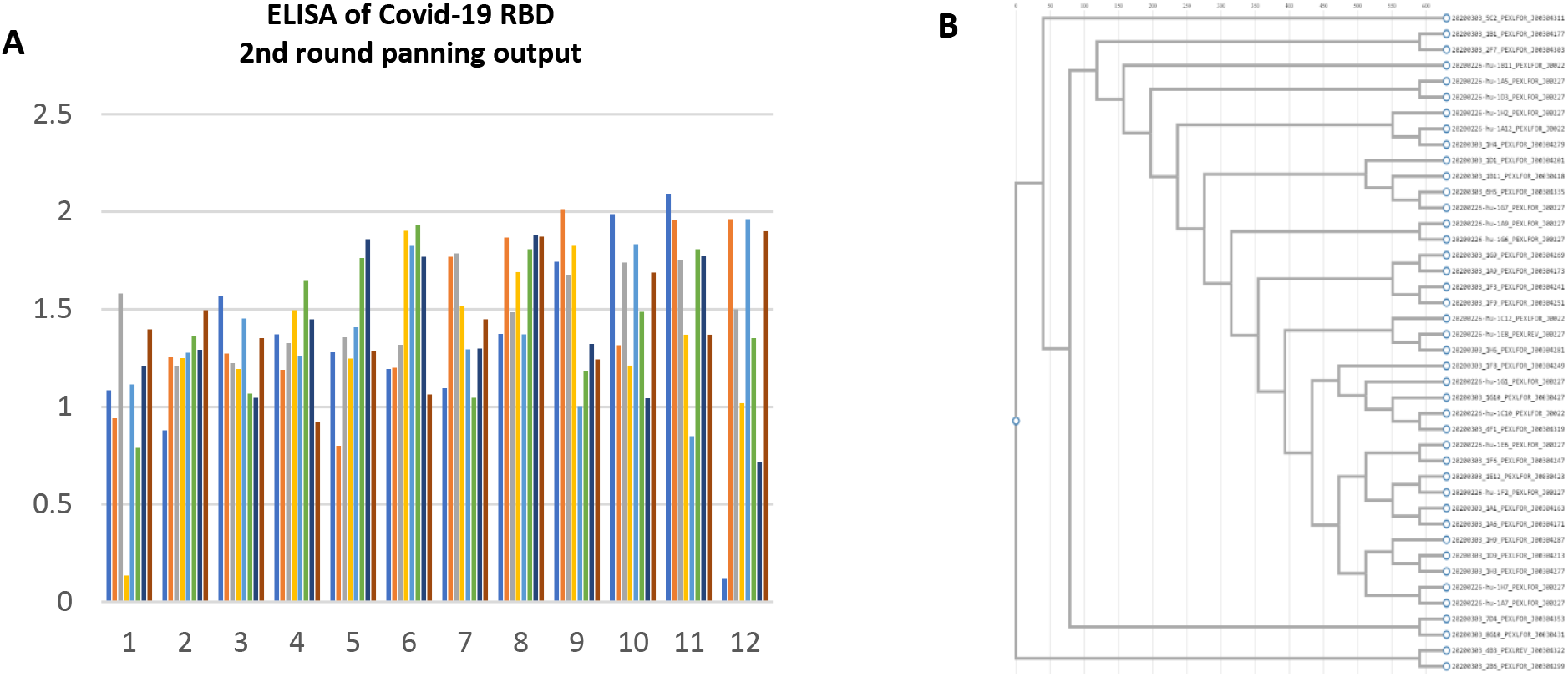
Monophage ELISA screening Covid-19 RBD phage antibodies. EHL library was panned with Covid-19 for 2 rounds. Single colonies form 2^nd^-round output was picked to prepare monoclonal phage antibody solutions, examined by ELISA with RBD, mouse Fc and 293F cells. Only RBD positive hits were resorted to DNA sequencing of scFv genes. **A**. Chart of absorbance (A450) on RBD plate ELISA. **B**. Amino acid sequence multi-alignment of unique scFv clones positive for RBD binding.

By checking the amino acid sequences, we saw majority of the hit clones (37/42) belong to lambda scFv and only five belong to kappa scFv (data not shown), though equal amount of kappa and lambda scFv library aliquots were mixed before first panning. Among the lambda scFv hits, most of them (30/37) have almost same complementarity-determining region (CDR) compositions in both VH and Vλ, with sporadic differences in the framework region (FR), implying fast enrichment of antibody recognizing certain epitope.

### Membrane spike bound antibodies are specific for Covid-19

Phage antibody hits tend to be false positive by ELISA detection, due to the stickiness of phage proteins^18^ and epitope/conformation differences between purified/truncated protein and membrane/full-length parental molecule^19^ or lose the reactivity because the once exposed epitopes in protein fragment become inaccessible in native protein. This is very true because in current case, the panning bait is a dimeric and fragmented RBD (RBD-mFc), while Covid-19 native spike is a trimer^20^. To verify the authenticity of the phage antibody binders and check their ability to recognize cell membrane trimeric Covid-19 Spike, we first generated soluble antibodies to facilitate further characterization of their properties. Since many clones are highly similar in CDR residues and only differ by a few FR residues, to reduce the work we transferred 22 scFv genes (one of every two close members from above alignment) from phagemid to an in-house mammalian expression vector pFP and expressed them as scFv-huFc (human Fc) fusion. Upon protein-A purification and aggregates removal to make sure monomer achieved over 95%, the antibodies were tested by FACS on above-generated ID8 cells to screen positive membrane covid-19 spike binders, which are the potential neutralizing antibody candidates.

8 out of the 22 expressed scFv-huFc antibodies demonstrated significant ID8 cell-binding capabilities with varied shifts by FACS (Fig.3). We tried to rank the binders roughly by the positive percentages, which is the reflection of apparent affinity instead of intrinsic affinity. Clone 1B1 seems to be the strongest spike binder among them. 1B11 and 5C2 are comparable intermediate ones and the rest five are close as mild ones. Reviewing their amino acid sequences (data not shown), we noticed that 1B11 share almost the same CDR compositions are the rest five hits except 5C2 and 1B1but differs in several FR residues, however 1B11 exhibited significant higher P2 than its cognate members, indicating affinity beneficial effect of those different FR residues in 1B11. Future kinetics measurement should reveal the quantitative affinity gap.

**Fig. 3.**
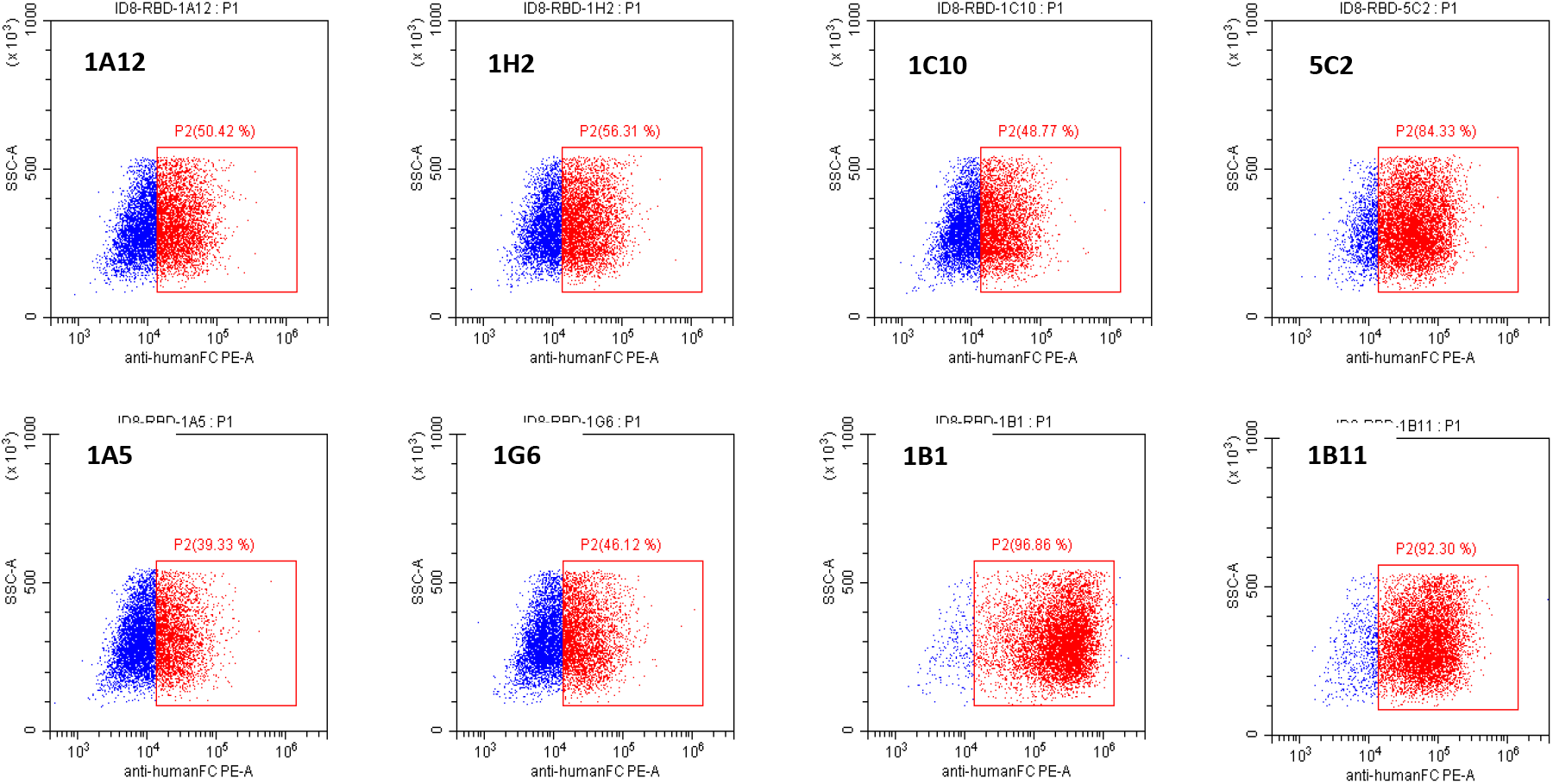
FACS examination of RBD-binding antibodies to Covid-19 membrane spike on ID8 cells. Soluble scFv-huFc of 22 hits that were RBD ELISA-positive were incubated with equal amount of ID8 cells, followed by goat anti-human Fc (PE conjugated). Stained cells were analyzed to draw dot-plot by FACS. P2 Gating was set based on the background of secondary antibody staining and considered positive. Only clones having positive percentages were shown. Individual clone was labeled in each plot.

It is now known that the amino acid sequence homologies between MERS-CoV, SARS-CoV-1 and SARS-CoV-2 are very high, especially the latter two, approximates to 75% for the spike proteins and are 73.7% for RBDs^3^. More conserved in structure than primary sequence, the RBDs in both viruses bind the same ACE2 in different affinities^21^. Genetically it has been revealed that in RBD, some area is conserved, and some are hypervariable^22^. To investigate whether the 8 ID8 cell-binding positive antibodies from above have any cross reaction to closely related spike proteins of SARS and MERS coronavirus, they were further examined by ELISA across SARS spike protein (SARS-S), MERS Spike protein (MERS-S) and Covid-19 spike protein (Covid-19-S). As it showed in Fig.4, All 8 hits are exclusively specific for Covid-19 spike and none of them has any cross reactivity to the other two spikes, indicating structurally conserved RBDs are not quite immunogenic conservative among these coronaviruses.

**Fig. 4.**
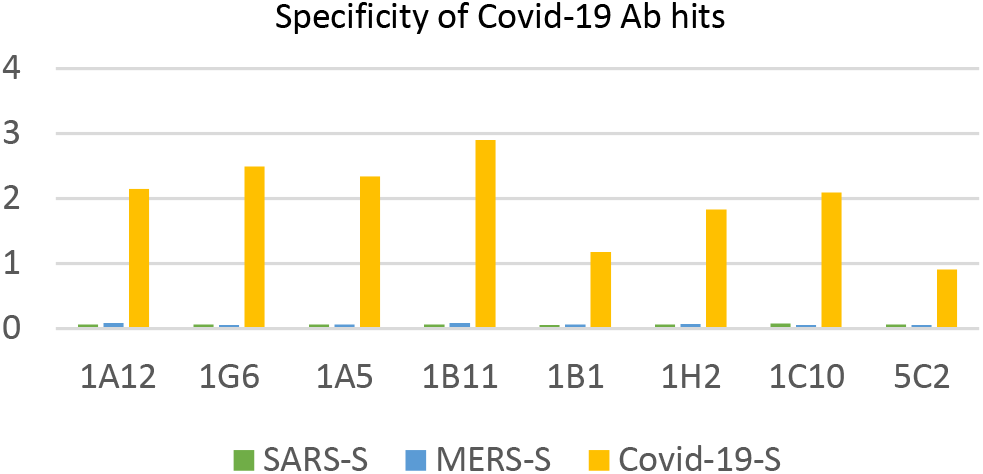
Cross-reaction examination of Covid-19 RBD hits by ELISA. Binding of 8 COVid-19 RBD-hits was examined by ELISA to the spike proteins (named as SARS-S, MERS-S and Covid-19-S) for SARS-COV-1, MERS-COV and SARS-COV-2 (Covid-19) and measured by absorbance (A450). All hits demonstrated strong positive to Covid-19-S (yellow bars) and none cross-reacted to the other two spikes (green and blue bars).

### The Covid-19 RBD specific antibodies bind to different epitopes

Having obtained 8 Covid-19 spike specific human antibodies, we wondered how their binding epitopes locating in its RBD and, more importantly, what is the potential impact on the binding of RBD to ACE2. Biolayer Interferometry (BLI) based instrument Blitz offers a simple straight forward evaluation method of antibody-receptor interaction at protein level^23^. Antibodies, RBD or ACE2 were loaded to appropriate sensors sequentially, typically that the previous step has been well equilibrated. The ascending or flat response curves recorded corresponding component incubated with sensors implied yes/no interaction among proteins between adjacent layers (Fig.5). Some observations are clear to see from the Blitz results: All hit clones bind to RBD by Blitz (Fig.5, A,B,C); 1B1 can concurrently bind to RBD with the other 6 antibodies (such as 1B11 etc.) except 5C2 (Fig.5 B,D); 1B1 mutually competes with 5C2 (Fig.5, C). Consistent with the competition between 1B1 and 5C2, when RBD is bound by 1B1, it can no longer be bound by 5C2, and vice versa. (Fig.5, D).

**Fig. 5.**
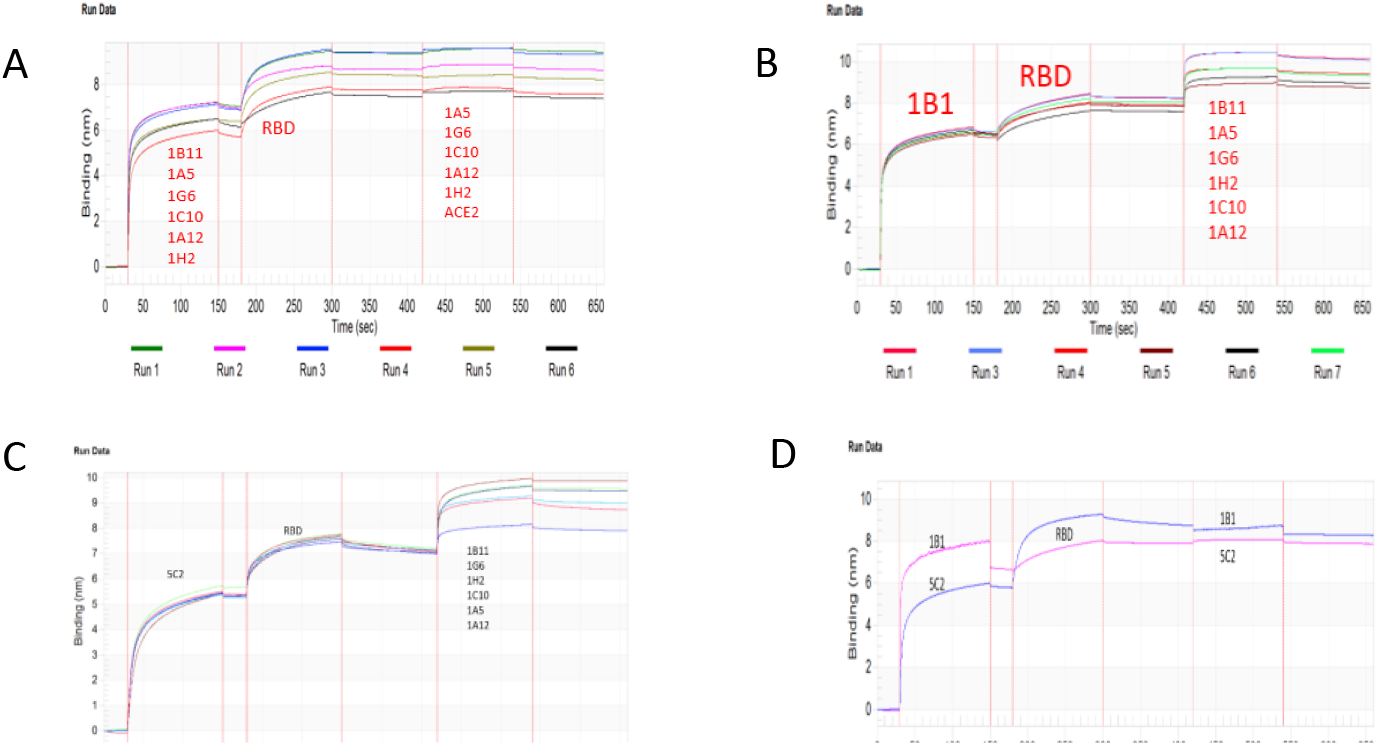
BLITZ epitope mapping of Covid-19 RBD-hits antibodies. (A). Rotational mutual interaction examination of the six hits (1B11, 1A5, 1G6, 1C10, 1A12 and 1H2) was captured by protein A sensors, followed by RBD binding and one of the six antibodies plus ACE2. RBD was bound well by individual antibodies but not concurrent binding. (B). 1B1 concurrently bound to RBD with any one of the six other antibodies (1B11, 1A5, 1G6, 1C10, 1A12 and 1H2). (C). 5C2 concurrently bound to RBD with any one of the six other antibodies (1B11, 1A5, 1G6, 1C10, A12 and 1H2). (D). 1B1 and 5C2 mutually competed in binding to RBD

Combining the amino acid sequence information and this BLI study, we drew a putative Venn epitope map of the 8 RBD-antibodies and ACE2 on Covid-19 RBD (Fig.6). For the convenience of description, we categorized the epitope as group I (1B1 antibody), group II (5C2 antibody) and group III (antibody 1B11, 1A5, 1A12, 1G6, 1H2 and 1C10).

**Fig. 6.**
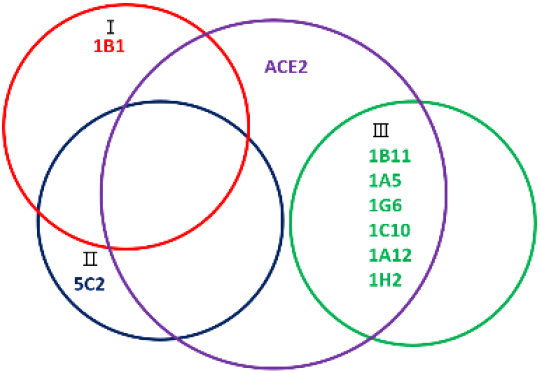
Venn map of Covid-19 hits and ACE2 on RBD. Epitope locations on RBD by the isolated 8 hit antibodies were deduced from the above Blitz results and largely categorized into 3 groups (I, II and III). 1B1 and 5C2 have large overlapped sites in RBD and may block the most part of RBD-ACE2 interaction interface. The group III hits (1B11, 1A5, 1G6, 1C10, 1A12 and 1H2) binds to an independent site from 1B1 or 5C2, however may block a minor side of ACE2-RBD interaction interface.

### Spike+ cells are bound simultaneously by RBD-antibodies and ACE2

In silicon examined by Blitz done above on the obtained RBD-antibodies implied their potential varied interferences to the ACE2 interaction with Covid-19 Spike. Since their epitopes/affinities are different, we started to investigate whether competition exists, or how much is it, between these antibodies and ACE2 in binding to ID8 cells. A preliminary experiment was done to check if there was dual binding when both ACE2 and individual antibodies were added to ID8 cells. Equal concentration of ACE2-mFc fusion (mouse Fc, MW 110KD) and any one of the 8 RBD-specific scFv-huFc (human Fc, MW 100KD) were mixed and incubated with ID8 cells before appropriate 2^nd^ fluorescent reagents were added for FACS analysis. Dot-plots of PE and FITC channels were depicted (**Fig.7**) for every sample.

**Fig. 7.**
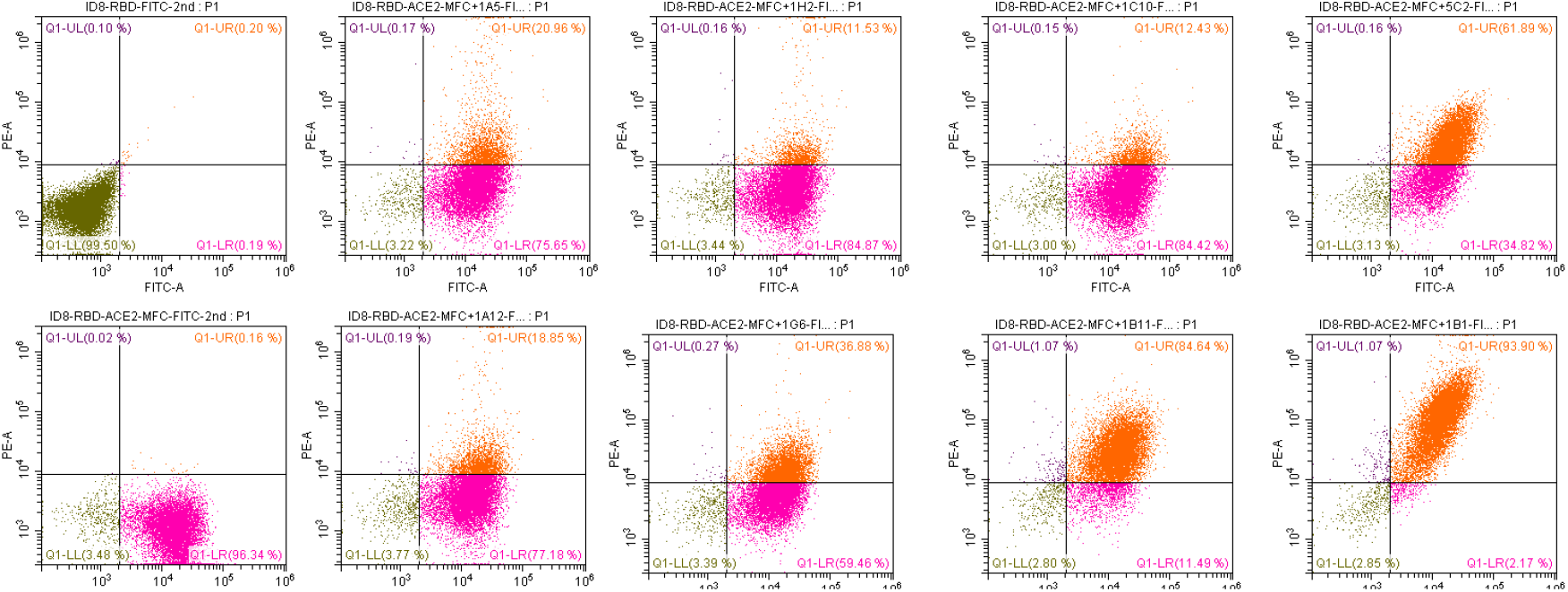
Dual binding examination of Covid-19 RBD-hit antibodies. All 8 RBD hits separately were incubated together and ACE2-mFc at around equal concentrations before adding to ID8 cells. Both antibody binding and ACE2 binding were detected by corresponding specific secondary antibodies (different fluorescent conjugation). Upper row from left: FITC-conjugated secondary control, 1A5, 1H2, 1C10, 5C2. Lower row from left: ACE2-mFc (alone, positive control), 1A12, 1G6, 1B11 and 1B1

Dot-plots of all groups demonstrated strong ACE2 binding (FITC channel), which is consistent with earlier study that ID8 expresses functional Covid-19 Spike. On one side, the fact that ACE2+ population in all groups are greater than 95% (Fig.7) implied limited blockade of RBD-antibodies to prevent Covid-19 spike from binding to ACE2 when both are present simultaneously at around equal concentration. On the other side, all antibodies demonstrated positive binding to the ID8 cells as well, at varied extent from~11% (1H2) to ~94% (1B1). Again, 1B1 championed the ranking, followed by 1B11 and 5C2. By the percentage of dual positive population, we can easily rank the antibodies in order of 1B1>1B11>5C2>1G6 etc. for their competitiveness against ACE2. Surprisingly the rest four antibodies (1H2, 1C10, 1A5 and 1A12), who share the same CDR compositions (but not exact amino acids in FR) as 1B11 and 1G6 (thus the similar/same epitope), showed significant less binding on cells, implying weaker affinities caused by the different FR residues. Overall the data here proved that there is varied extent of competitiveness between these RBD antibodies and ACE2 in binding to Covid-19 spike.

### RBD nAbs preventively block interaction between Covid-19 spike and ACE2

Next we explored another scenario where antibodies are much abundant than ACE2 (this is possible since antibody can be administered beforehand in large amount as prophylaxis measure). In this case we wanted to find out if any of these RBD-hits can significantly reduce the binding of spike to ACE2, i.e., neutralize the virus (here again we used Covid-19 spike-expressing ID8 as surrogate). We first titrated the concentration of ACE2-mFc on fixed amount of ID8 cells. ACE2 is abundantly and widely expressed in human epithelial cells, the total (the soluble and membrane) expression level is yet to know, although ACE2 in the serum level has been reported^24^. We found that when ACE2-mFc was around 0.02 μg/ml, ~100% ID8 cells were positive by FACS (Fig.8, green histograms), indicating very strong interaction between Covid-19 spike and ACE2. For the neutralization assay, we set up a serial dilution (2 times down) of every candidate antibody from the 10 μg/ml to ~0.31 μg/ml (a point where there was minimal impact on ACE2-mFc binding to spike+ cells in a preparatory experiment), added them individually to same fixed amount of ID8 cells and incubated enough time to saturate the cells (spike). Next, the saturation concentration of ACE2-mFc was loaded to compete out antibody. Finally, only signals from FITC channel was collected, which were the MFI (mean fluorescence intensity) numbers of ACE2-mFc binding to ID8 cells. Histograms of part of the diluted sample concentrations (shown in Fig.8) clearly demonstrated that some the RBD-hits, such as 5C2, can objectively block Covid-19 spike from binding to ACE2 at various concentrations.

**Fig. 8.**
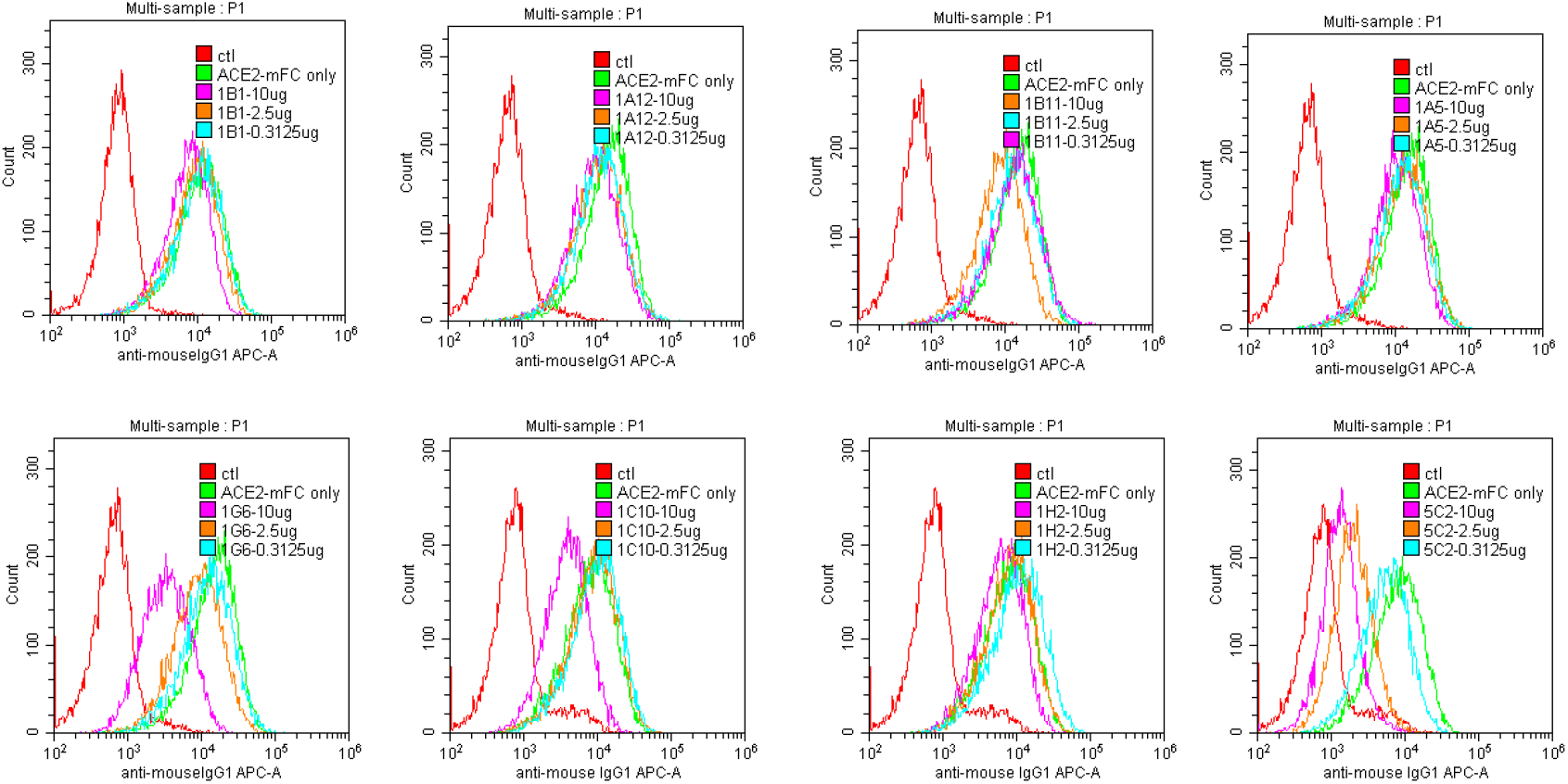
Competition FACS evaluation of neutralizing potentials of RBD-hit antibodies. Serial diluted RBD antibodies were preincubated separately with ID8 cells before adding of ACE2-mF c, whose binding was analyzed by APC-conjugated secondary antibody. The FACS plot of ACE2-mFc with or without antibody is shown. Negative control (red), no antibody ACE2-mF c control (green), and only histograms of three antibody concentrations 10 μg/ml plots (pink), 2.5 μg/ml plots (yellow) and 0.31 μg/ml plots (aqua) were shown. Upper row from left: 1B1, 1A12, 1B11, 1A5. Lower row from left: 1G6, 1C10, 1H2 and 5C2.

When there is competition between any RBD hit antibody and ACE2 in binding to Covid-19 spike, an increased antibody concentration would lead to reduced ACE2 attachment, quantitatively represented by MFI. Indeed, after grouping the MFIs as shown in Fig.9, at 10 μg/ml (a clinically achievable serum concentration) we observed among 8 of the antibodies, at least 3 hit antibodies (1G6, 1C10 and 5C2) demonstrated significant ACE2 attachment reduction, decreasing the MFI of ACE2-mFc by as much as 60%, in comparing to negative antibody control. The rest 5 hits caused little to moderate inhibition at this concentration. As antibody concentration lowered down (less antibody binding), MFI increased, more ACE2-mFc occupied the ID8 cells. Overall a dose-dependent manner is observed for the antibody inhibition. Among them, 5C2 performed the best in neutralizing Covid-19 spike protein in binding to ACE2-mFc across all dilutions to the lowest point, with around 50% reduction of MFI. Surprisingly, contrary to the previous ranks where 1B1 seemed to be best in binding to RBD and probably have large portion of overlapped epitopes with 5C2, it demonstrated much mild inhibition of spike in binding to ACE2 in this assay. This finding highlighted the importance of epitope location in order to be an efficient blocker besides decent affinity.

**Fig. 9.**
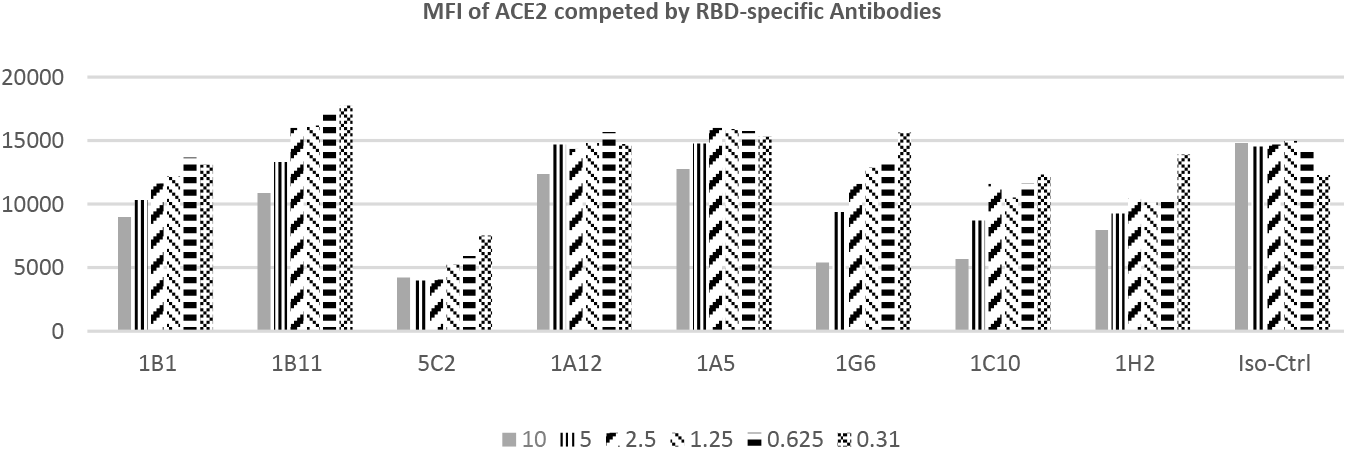
MFI change of ACE2 binding to ID8 cells in the presence of serial diluted RBD-antibodies. 8 Covid-19 spike positive antibody binders (1B1, 1B11, 5C2, 1A12, 1A5, 1G6, 1C10 and 1H2) and one isotype negative control antibody (iso-ctrl in the figure) were mixed with ID8 cells at serial diluted concentrations (10 μg/ml down to 0.31 μg/ml, 2 times dilution) before adding of human ACE2-mFc, which has strong interaction with ID8. MFIs (shown in vertical axis) of individual antibody/concentrations cell group (horizonal axis) were complied.

## Discussions

In the report we described the construction of a naïve human antibody phage display library and the usage of it in isolation meaningful weapons against Covid-19. We have preliminarily characterized the epitopes, specificity and neutralization potentials of the obtained human antibodies to Covid-19 spike. Our study offered promising human nAb candidates for further development of antibody-based weapons against the SARS-COV-2.

Antibody Phage display technology has contributed antibody drugs such as FDA-approved blockbuster (Humira) and many clinical stage testing candidates^25^. Generally, immune libraries are preferred for the generation of antibodies against infectious disease related targets due to the biased antibody repertoires as a result of exposure and in vivo evolution of immune response to the infection^26^. However, the target specific libraries must be constructed each time to get enriched antibodies against antigens from different infections. A human PBMC-derived naïve phage naïve display antibody library is a ready-to-go tool, being particularly valuable resources to general targets and shortened R&D process when developing antibody-based clinical therapeutics. Diversity and library size are the two most critical parameters in generating a useful library for any given target^27^. In the study we first reported how such a library was built and became a key resource to the R&D of antibody-based diagnostics and biotherapeutics.

With that advantage we quickly launched a campaign to screen hits, targeting Covid-19 spike (S) protein. Since the outbreak of SARS, many reports have published to elucidate how the spike of corona virus is responsible for interacting with host cell receptors, mainly human ACE2 to initiate infection^28, 29^. Structurally, the receptor binding domain (RBD) locating from AA 321-535 in S protein, is the exact portion that interacts with ACE2^12, 22^. Aiming to have better chance in obtain potent nAbs that can efficiently block RBD-ACE2 interaction, we used engineering expressed RBD as bait in solid phase biopanning. Dozens of unique primary positive phage antibodies to RBD were selected by monophage ELISA and proved to be very specific for Covid-19, with no cross binding to the other two closely related corona virus members, SARS-CoV-1 and MERS-CoV. Through BLI analysis on Blitz these antibodies span at least two independent groups of epitopes, each occupying part if not the complete ACE2 binding site in RBD. Additionally, their viral neutralizing potency was evaluated by competition FACS. 3 of the 8 hits demonstrated mild to significant competition in a concentration dependent mode to reduce spike binding to ACE2, suggesting their preventative and therapeutic potentials in combating the Covid-19 pandemic. One of the hits championed the competition assay, exhibited promising development value. Preliminary microneutralization study on Covid-19 pseudo virus has proved some of the RBD-antibodies in this report blocked viral infection (raw data).

Since outbroke last December to this date the Covid-19 viral destruction is still ongoing with no foreseeable end. Without any validated medicines clinically available currently, diagnostic, preventive and therapeutic remedies are desperately needed should the virus last long and come back soon. NAbs are historically effective in fighting against viral pandemics as they effectively inhibit virus’s entry by preventing viral attachment or membrane fusion^30, 31^ and very likely to play a critical role to fight Covid-19^32^. As late as from the outbreaking of SARS, MERS, Ebola to Zika, to Covid-19 convalescent plasma^7, 33, 34^ which may contain polyclonal nAbs collected from recovered patients have been tested to treat seriously ill people and have shown certain reduction protection in some cases. Scientific studies of the nAbs to SARS, MERS on animal models proved their protection effect. Currently researchers all over the world are racing to isolated nAbs from Covid-19 immunized animals (llama)^35^, available SAR-Cov-1 neutralizing antibody^36^ and/or human B cells of infected patients^37^ to develop potential therapeutics.

Classical antibody neutralization is strictly defined as the reduction in viral infectivity by the binding of antibodies to the surface of viral particles (virions), thereby blocking a step in the viral replication cycle that precedes virally encoded transcription or synthesis^38, 39^. In that regard nAbs are the best correlate of protection from viral infection after vaccination. Nabs offers the direct function of abolition of a pathogen’s infectivity upon binding, with no participation of any other components of the innate or adaptive immune system. So, neutralization is probably the most robust and powerful function that antibodies exert against viruses. In a broadly sense antibodies can neutralize viral infectivity in several additional ways^40, 41^. They may block viral uptake into cells, prevent uncoating of the genomes in endosomes, cause aggregation of virus particles, or lyse the viral membrane through by antibody-dependent cellular cytotoxicity (ADCC) and antibody dependent cellular phagocytosis (ADCP). These protective antibodies are targeting components in other viral cycle steps, as well as utilizing human immune system to clear virus^42^.

While there is little doubt that clinical Covid-19 nAbs can be developed sooner or later, there is a big hurdle in front of us to overcome, that is, strain coverage and escape. Diversities of collected 86 variant covid-19 genomes have been recently unveiled^43^. Of note, deletions in spike glycoprotein and mutations located in RBD have been found. More mutations are likely to emerge as the virus continues to transmit across ethics and global regions. Subsequently monoclonal antibody reacting to specific epitope in RBD is probably vulnerable to the mutations and glycosylation modifications. Additional studies to dissect the exact recognizing residues of any candidate neutralizing Covid-19 antibody are necessary before moving to further development. Alternatively, as backup it’s rational to develop broad nAbs to more conservative regions in spike^44, 45^. In fact, nAbs targeting a highly conserved epitope in RBD has been isolated from convalescent plasma of SARS patient^46^. Combination therapy with multiple non-competing nAbs recognizing different epitopes on the RBD/spike may be another ideal direction to avoid immune escape and increase coverage^47^.

In summary, we have reported here the isolation and characterization of nAbs to Covid-19 RBD from a human naïve phage display library. Our work provided promising antibody candidates as potential diagnostic, prophylaxis and therapeutic reagents for further development.

## Materials and Methods

### Construction of EHL human naïve phage display library

A very large naïve human single chain variable fragment (scFv) phage display library was constructed similarly as previously reported^15^. Briefly, PBMCs (10 million/donor, purchased from BuyPBMCs, Precision for Medicine, https://buypbmcs.com/) collected from healthy donors to extract the total RNA (Cat#74104, RNeasy^®^, QIAGEN, Valencia, CA). First strand cDNA was generated by using SuperScript™ III First-Strand Synthesis System (ThermofisherScientific, cat#18080051) from every RNA sample (5 μg RNA /rxn) with random hexamer primers or Anchored Oligo(dT)20 (ThermofisherScientific, cat#12577011) before pooling. Variable heavy (VH) and light (Vκ and Vλ) genes were amplified separately from the pooled cDNA samples by two rounds of PCR using HotStarTaq (Qiagen, Cat# 203443) and human antibody sequences primers derived from previous report^15^. A pair of SfiI sites were incorporated into the 5’-end of VH primers and 3’-end of Vκ and Vλ primers to facilitate followed phagemid cloning. Overlapping of VH-Vκ or VH-Vλ scFv genes were assembled separately. Amplified scFv PCR fragments were purified with Gel Extraction kit (Qiaquick Gel Extraction Kit, Cat# 28706, QIAGEN), and subjected to SfiI (Fast Digest SfiI, ThermofisherScientific, cat# 1824) digestion.

To construct the scFv phagemid library, vector pADL-10b (Antibody Design laboratories, cat#PD0105) was completely digested with SfiI (ThermoFisherScientific, cat#FD1824) followed by dephosphorylation treatment (CIAP, ThermoFisherScientific, cat#18009027) before purification with PCR purification kit (QIAquick PCR Purification Kit, cat#28104). Digested scFv and pADL-10b was ligated by the ratio of 3:1, using T4 DNA ligase (NEB, Cat#M0202) and incubation overnight at 12°C. Ligation mixture was purified and electroporated into TG1 E. coli (Lucigen, Cat# 60502-2) over fifty shocks for each library. Combined TG1 slurry culture was grown in non-expression (NE) media (2YT containing 1% glucose and 100 μg/ml carbenicillin) at 37°C for one hour before a serial of 10 times dilutions were made to measure the approximate size of the kappa or lambda scFv library. The rest culture was expanded in NE media, grew to OD600 around 2, and aliquots of culture were frozen as seeds for future production of phage libraries. Propagation of combinatorial phage libraries rescued by M13K07 helper phage (Antibody Design laboratories, cat#PH010L) was performed as published procedures^47^. Phage antibody library aliquots were stored at −70°C in a solution of 50% glycerol and named as EHL library.

### Construction of Covid-19 spike-expressing cell line

To express the spike (S) protein of SARS-CoV-2 in mammalian cells, a codon optimized cDNA (GenBank:NC_045512.2) encoding the full-length S protein, a transmembrane motif (TM) and 3xFLAG tag was synthesized and cloned into a pRRL-derived vector (R48) by XbaI/XmaI, yielding pRRL-19S-FLAG-BSD. 293FT cells were transfected with lenti-viral made from SARS-CoV-2 S plasmid and other vectors (pLP-1-Gag/Pol, pLP-2-Rev and pLP-VSVG, all in-house made) containing packaging elements. Media was replaced once 12-16 hours after transfection. Pseudo virus stock (named as LV-19S) was collected from the supernatant and filtered through 0.45 μm membrane.

To build stable cell line expressing Covid-19 S, ID8 cells (a gift from Institute of immunotherapy, Fujian Medical University) was seeded at 10,000/well in DMEM (Gibco, cat#31600-034) +10% FBS (GeminiBio, cat#100-500) in6-well culture plate and incubated overnight. The next morning, 1ml of LV-19S (and vehicle control) was added and incubation continued 24 hours before changing the media with Screening media (DMEM+10% FBS containing 10 μg/ml Blasticidin S hydrochloride (Invitrogen, cat#21001)). Screening media was changed every 3 days until day 7 when non-infected cells all gone. Continue the infected cell culture, which is the stable cell line, maintained in DMEM+10% FBS containing 8 μg/ml Blasticidin S hydrochloride.

### Panning of RBD and monophage ELISA

Solid phase biopanning of hits from a phage display antibody library was conducted as previously reported^49^ with minor modifications in library blocking. Briefly Covid-19 spike RBD (Sinobiological, cat#40592-V05H, mouse Fc fusion) protein was coated at 2μg/ml in wells of a 96-well Nunc Maxisorp microplate and placed in a refrigerator overnight. One each of Kappa and lambda EHL Library aliquots were mixed and blocked with 4% milk PBS (MPBS), 10 million 293F cells (to deplete potential binders to residual protein contaminated in vendor’s RBD) and 10μg/ml mouse IgG (to deplete mouse Fc binders) before adding the library stock to the RBD-coating wells. The rest steps are standard protocol^49^. Two to three cycles of biopanning were performed before ELISA screening was conducted to select the specific RBD-binding phage antibodies from the output libraries. Monophage ELISA was carried out essentially as described^50, 51^ with RBD as coating antigen, mouse Fc and 293F cells as negative control.

### Soluble expression and purification of scFv-Fc

Unique scFv clones from DNA sequencing analysis were selected for further subcloning and fusion expression with human Fc. ScFv genes were cut and paste with SfiI and NcoI sequentially from the phagemid vector into pFP vector (in-house made). Freestyle 293F (ThermoFisherScientific, Cat#R79007) cell was transfected with purified miniprep plasmid DNA (QIAgen minipreps kit, Qiagen, Cat#29107) as described (Lipofectamine™ RNAiMAX Transfection Reagent, ThermoFisher Scientific, Cat#13778500) in 30-ml volume. scFv-Fc product was purified by protein-A affinity Purification column (HiTrap^®^ Protein A High Performance, GE Life science, cat#GE17-0402-01) and size exclusion chromatography (GE SEC, superdex 200, GE Life science, cat#GE17-0612-10) to remove aggregates, whenever necessary. Monomer percentage was analyzed by HPLC-SEC^52^.

### Blitz characterization of hits and epitope mapping

The binding sites of obtained antibodies to RBD was examined by Blitz system (ForteBio, Pall Lifesciences). 10 μg/ml Biotinylated RBD protein was captured by streptavidin biosensor (ForteBio, cat# 18-5019), followed by sequential binding of 10 μg/ml RBD antibody A then 10 μg/ml antibody B, in the presence of 10 μg/ml antibody A, or vice versa. Alternatively, 10 μg/ml antibody A was captured by anti-human Fc biosensor (ForteBio, cat# 18-5060) until saturation, followed by sequential binding of 10 μg/ml RBD, antibody B. Interaction curves were collected and analyzed to determine the relative binding locations of the antibodies on RBD.

### FACS examination of hits binding to Spike

Spike expressing cell 1D8 was maintained in 1640 media. To examine if spike trimer was expressed, both FITC conjugated anti-FLAG (CST, cat#8146) and ACE2-mFc (Sinobiologics, cat#10108-H05H) and FITC-anti-mouse-IgG (Biolegend, cat#405305) were incubated with 1-2 million 1D8 cells following standard FACS protocol and analyzed by Beckman.

To examine if the antibody hits from ELISA screening were still capable of binding to native spike trimer, the individual antibody hit at 5 μg/ml and/or ACE2-mFc was incubated with 1-2 million 1D8 spike+ cells and detected by anti-human-Fc PE (invitrogen, 12-4998-82) and FITC Goat anti-mouse-IgG (Biolegend, cat#405305) or APC Goat anti-mouse-IgG (Biolegend, cat# 405308) following standard FACS protocol and analyzed by Beckman FACS (CytoFLEX, CytExpert2.0).

### Neutralization assay of RBD-antibodies by FACS

To check whether ACE2-Spike interaction can be blocked by the isolated nAbs, individual titrated antibody from 10 ug/ml down to 0.31 ug/ml (2 times down) was incubated with 1-2 spike+ cells for 1 hour at 4°C before adding of 0.02 ug/ml ACE2-mFc followed by FITC conjugated anti-mouse IgG (Biolegend, cat#405305) and the MFI of FITC and positive percentage were collected.

### Statement

Andy Q. Yuan is the co-founder and CSO of Yikesite Biopharma Development LLC, which wholly-owns Excyte LLC. Qingwu Meng is the co-founder of Yikesite Biopharma Development LLC. Likun Zhao, Lili Bai, Yanhu Li, Daqing Guo, Shanshan Zhen, Xiaojun Chen, Ji Yang and Xiaoying Xue are the employees of Yikesite Biopharma Development LLC.

## Notes

### Competing Interest Statement

Andy Yuan is the co-founder and CSO of Yikesite Biopharma Development LLC, which wholly-owns Excyte LLC. Qingwu Meng is the co-founder of Yikesite Biopharma Development LLC. Likun Zhao, Lili Bai, Yanhu Li, Daqing Guo, Shanshan Zhen, Xiaojun Chen，Ji Yang and Xiaoying Xue are the employees of Yikesite Biopharma Development LLC.

## References

1. https://www.worldometers.info/coronavirus/

2. S.K. Lal (ed.), Molecular Biology of the SARS-Coronavirus, DOI 10.1007/978-3-642-03683-5_1, ©Springer-Verlag Berlin Heidelberg 2010

3. Chunyun Sun, Long Chen, Ji Yang, Chunxia Luo, Yanjing Zhang, Jing Li, Jiahui Yang, Jie Zhang, Liangzhi Xie. SARS-CoV-2 and SARS-CoV Spike-RBD Structure and Receptor Binding Comparison and Potential Implications on Neutralizing Antibody and Vaccine Development doi: https://doi.org/10.1101/2020.02.16.951723

4. Hoffmann et al., 2020, Cell 181, 271–280 April 16, 2020 ^a^ 2020 Elsevier Inc. https://doi.org/10.1016/j.cell.2020.02.052

5. Driver LC, Oertel MD. Synagis: an anti-RSV monoclonal antibody. Pediatr Nurs. 1999 Sep-Oct;25(5):527–30.

6. Romero JR. Palivizumab prophylaxis of respiratory syncytial virus disease from 1998 to 2002: results from four years of palivizumab usage. Pediatr Infect Dis J. 2003 Feb;22(2 Suppl):S46–54.

7. Rajendran K, Narayanasamy K, Rangarajan J, Rathinam J, Natarajan M, Ramachandran A. Convalescent plasma transfusion for the treatment of CoVID-19: Systematic review. J Med Virol. 2020 May 1. doi: 10.1002/jmv.25961. Review.

8 Coughlin MM, Prabhakar BS. Neutralizing human monoclonal antibodies to severe acute respiratory syndrome coronavirus: target, mechanism of action, and therapeutic potential. Rev Med Virol. 2012 Jan;22(1):2–17. doi: 10.1002/rmv.706. Epub 2011 Sep 9. Review.

9. Han HJ, Liu JW, Yu H, Yu XJ. Neutralizing Monoclonal Antibodies as Promising Therapeutics against Middle East Respiratory Syndrome Coronavirus Infection. Viruses. 2018 Nov 30;10(12). pii: E680. doi: 10.3390/v10120680. Review.

10. Li, W., M. J. Moore, N. Vasilieva, J. Sui, S. K. Wong, M. A. Berne, M. Somasundaran, J. L. Sullivan, K. Luzuriaga, T. C. Greenough, H. Choe, and M. Farzan. 2003. Angiotensin-converting enzyme 2 is a functional receptor for the SARS coronavirus. Nature 426:450–454

11. Tai W, He L, Zhang X, Pu J, Voronin D, Jiang S, Zhou Y, Du L. Characterization of the receptor-binding domain (RBD) of 2019 novel coronavirus: implication for development of RBD protein as a viral attachment inhibitor and vaccine. Cell Mol Immunol. 2020 Mar 19. doi: 10.1038/s41423-020-0400-4.

12. Yan R, Zhang Y, Li Y, Xia L, Guo Y, Zhou Q. Structural basis for the recognition of SARS-CoV-2 by full-length human ACE2. Science. 2020 Mar 27;367(6485):1444–1448. doi: 10.1126/science.abb2762. Epub 2020 Mar 4.

13. Marks JD, Hoogenboom HR, Bonnert TP, McCafferty J, Griffiths AD, Winter G. By-passing immunization. Human antibodies from V-gene libraries displayed on phage. J Mol Biol. 1991 Dec 5;222(3):581–97.

14. Clackson T, Hoogenboom HR, Griffiths AD, Winter G. Making antibody fragments using phage display libraries. Nature. 1991 Aug 15;352(6336):624–8.

15. Yuan, Q., Robinson, M.K., Simmons, H.H. et al. Isolation of anti-MISIIR scFv molecules from a phage display library by cell sorter biopanning. Cancer Immunol Immunother 57, 367–378 (2008). https://doi.org/10.1007/s00262-007-0376-2

16. Wrapp D, Wang N, Corbett KS, Goldsmith JA, Hsieh CL, Abiona O, Graham BS, McLellan JS. Cryo-EM structure of the 2019-nCoV spike in the prefusion conformation. Science. 2020 Mar 13;367(6483):1260–1263. doi: 10.1126/science.abb2507.

17. Griffiths AD, Williams SC, Hartley O, Tomlinson IM, Waterhouse P, Crosby WL, Kontermann RE, Jones PT, Low NM, Allison TJ. Isolation of high affinity human antibodies directly from large synthetic repertoires. EMBO J. 1994;13:3245–3260

18. KK Murthy, I Ekiel, SH Shen, D Banville Fusion proteins could generate false positives in peptide phage display. BioTechniques 26:142–149

19. Sullivan N, Sun Y, Sattentau Q, et al. CD4-Induced conformational changes in the human immunodeficiency virus type 1 gp120 glycoprotein: consequences for virus entry and neutralization. J Virol. 1998;72(6):4694–4703.

20. Walls AC, Park YJ, Tortorici MA, Wall A, McGuire AT, Veesler D. Structure, Function, and Antigenicity of the SARS-CoV-2 Spike Glycoprotein. Cell. 2020;181(2):281–292.e6. doi:10.1016/j.cell.2020.02.058

21. Wan Y, Shang J, Graham R, Baric RS, Li F. Receptor Recognition by the Novel Coronavirus from Wuhan: an Analysis Based on Decade-Long Structural Studies of SARS Coronavirus. J Virol. 2020 Mar 17;94(7). pii: e00127–20. doi: 10.1128/JVI.00127-20.

22. Lan J, Ge J, Yu J, Shan S, Zhou H, Fan S, Zhang Q, Shi X, Wang Q, Zhang L, Wang X. Structure of the SARS-CoV-2 spike receptor-binding domain bound to the ACE2 receptor. Nature. 2020 Mar 30. doi: 10.1038/s41586-020-2180-5.

23. Kamat V, Rafique A. Designing binding kinetic assay on the bio-layer interferometry (BLI) biosensor to characterize antibody-antigen interactions. Anal Biochem. 2017 Nov 1;536:16–31. doi: 10.1016/j.ab.2017.08.002.

24. Tissue distribution of ACE2 protein, the functional receptor for SARS coronavirus. A first step in understanding SARS pathogenesis. Hamming I, Timens W, Bulthuis ML, Lely AT, Navis G, van Goor H. J Pathol. 2004 Jun;203(2):631–7.

25. Nixon AE, Sexton DJ, Ladner RC. Drugs derived from phage display: from candidate identification to clinical practice. MAbs. 2014;6(1):73–85. doi:10.4161/mabs.27240

26. Lim BN, Tye GJ, Choong YS, Ong EB, Ismail A, Lim TS. Principles and application of antibody libraries for infectious diseases. Review. Biotechnol Lett. 2014 Dec; 36(12):2381–92.

27. Sara Carmen and Lutz Jermutus. Concepts in antibody phage display. BRIEFINGS IN FUNCTIONAL GENOMICS AND PROTEOMICS. VOL 1. NO 2. 189–203. JULY 2002

28. Kuhn JH, Li W, Choe H, Farzan M. Angiotensin-converting enzyme 2: a functional receptor for SARS coronavirus. Cell Mol Life Sci. 2004 Nov;61(21):2738–43. Review.

29. Li F, Li W, Farzan M, Harrison SC. Structure of SARS coronavirus spike receptor-binding domain complexed with receptor. Science. 2005 Sep 16;309(5742):1864–8.

30. Coughlin MM, Prabhakar BS. Neutralizing human monoclonal antibodies to severe acute respiratory syndrome coronavirus: target, mechanism of action, and therapeutic potential. Rev Med Virol. 2012 Jan;22(1):2–17. doi: 10.1002/rmv.706. Epub 2011 Sep 8. Review.

31. Han HJ, Liu JW, Yu H, Yu XJ. Neutralizing Monoclonal Antibodies as Promising Therapeutics against Middle East Respiratory Syndrome Coronavirus Infection. Viruses. 2018 Nov 30;10(12). pii: E680. doi: 10.3390/v10120680. Review

32. Zhou G, Zhao Q. Perspectives on herapeutic neutralizing antibodies against the Novel Coronavirus SARS-CoV-2. Int J Biol Sci. 2020 Mar 15;16(10):1718–1723. doi: 10.7150/ijbs.45123. Review.

33. Mair-Jenkins J, Saavedra-Campos M, Baillie JK, Cleary P, Khaw FM, Lim WS, Makki S, Rooney KD, Nguyen-Van-Tam JS, Beck CR; Convalescent Plasma Study Group. The effectiveness of convalescent plasma and hyperimmune immunoglobulin for the treatment of severe acute respiratory infections of viral etiology: a systematic review and exploratory meta-analysis. J Infect Dis. 2015 Jan 1;211(1):80–90. doi: 10.1093/infdis/jiu396. Review.

34. Bloch EM, Shoham S, Casadevall A, Sachais BS, Shaz B, Winters JL, van Buskirk C, Grossman BJ, Joyner M, Henderson JP, Pekosz A, Lau B, Wesolowski A, Katz L, Shan H, Auwaerter PG, Thomas D, Sullivan DJ, Paneth N, Gehrie E, Spitalnik S,

35. Wrapp D, De Vlieger D, Corbett KS, Torres GM, Wang N, Van Breedam W, Roose K, van Schie L; VIB-CMB COVID-19 Response Team, Hoffmann M, Pöhlmann S, Graham BS, Callewaert N, Schepens B, Saelens X, McLellan JS. Structural Basis for Potent Neutralization of Betacoronaviruses by Single-Domain Camelid Antibodies. Cell. 2020 Apr 29. pii: S0092-8674(20)30494-3. doi: 10.1016/j.cell.2020.04.031.

36. Wang, C., Li, W., Drabek, D. et al. A human monoclonal antibody blocking SARS-CoV-2 infection. Nat Commun 11, 2251 (2020). https://doi.org/10.1038/s41467-020-16256-y

37. Fan Wu, Aojie Wang, Mei Liu, Qimin Wang, Jun Chen, Shuai Xia, Yun Ling, Yuling Zhang, Jingna Xun, Lu Lu, Shibo Jiang, Hongzhou Lu, Yumei Wen, Jinghe Huang Neutralizing antibody responses to SARS-CoV-2 in a COVID-19 recovered patient cohort and their implications. doi: https://doi.org/10.1101/2020.03.30.20047365

38. N. J. Dimmock, “Neutralization of animal viruses,” Current Topics in Microbiology and Immunology, vol. 183, pp. 1–149, 1993.

39. P. J. Klasse and Q. J. Sattentau, “Occupancy and mechanism in antibody-mediated neutralization of animal viruses,” Journal of General Virology, vol. 83, no. 9, pp. 2091–2108, 2002

40. Rhorer, J., Ambrose, C., Dickinson, S., Hamilton, H., Oleka, N., Malinoski, F., & Wittes, J. (2009). Efficacy of live attenuated influenza vaccine in children: A meta-analysis of nine randomized clinical trials Vaccine, 27 (7), 1101–1110 DOI: 10.1016/j.vaccine.2008.11.093

41. Saphire EO, Schendel SL, Gunn BM, Milligan JC, Alter G. Antibody-mediated protection against Ebola virus. Nat Immunol. 2018 Nov;19(11):1169–1178. doi: 10.1038/s41590-018-0233-9. Review.

42. Lewis GK, Pazgier M, Evans DT, Ferrari G, Bournazos S, Parsons MS, Bernard NF, Finzi A. Beyond Viral Neutralization. AIDS Res Hum Retroviruses. 2017 Aug;33(8):760–764. doi: 10.1089/AID.2016.0299. Review. Hod E, Pollack L, Nicholson WT, Pirofski LA, Bailey JA, Tobian AA. Deployment of convalescent plasma for the prevention and treatment of COVID-19. J Clin Invest. 2020 Apr 7. pii: 138745. doi: 10.1172/JCI138745. Review.

43. Phan T. Genetic diversity and evolution of SARS-CoV-2. Infect Genet Evol. 2020 Jul;81:104260. doi: 10.1016/j.meegid.2020.104260.

44. Preliminary Identification of Potential Vaccine Targets for the COVID-19 Coronavirus (SARS-CoV-2) Based on SARS-CoV Immunological Studies. Ahmed SF, Quadeer AA, McKay MR. Viruses. 2020 Feb 25;12(3). pii: E254. doi: 10.3390/v12030254.

45. Grifoni A, Sidney J, Zhang Y, Scheuermann RH, Peters B, Sette A. A Sequence Homology and Bioinformatic Approach Can Predict Candidate Targets for Immune Responses to SARS-CoV-2. Cell Host Microbe. 2020 Apr 8;27(4):671–680.e2. doi: 10.1016/j.chom.2020.03.002. Epub 2020 Mar 16.

46. Yuan M, Wu NC, Zhu X, Lee CD, So RTY, Lv H, Mok CKP, Wilson IA. A highly conserved cryptic epitope in the receptor-binding domains of SARS-CoV-2 and SARS-CoV. Science. 2020 Apr 3. pii: eabb7269. doi: 10.1126/science.abb7269.

47. Yan Wu, Feiran Wang, Chenguang Shen, Weiyu Peng, Delin Li, Cheng Zhao, Zhaohui Li, Shihua Li, Yuhai Bi, Yang Yang, Yuhuan Gong, Haixia Xiao, Zheng Fan, Shuguang Tan, Guizhen Wu, Wenjie Tan, Xuancheng Lu, Changfa Fan, Qihui Wang, Yingxia Liu, Jianxun Qi, George Fu Gao, Feng Gao, Lei Liu A non-competing pair of human neutralizing antibodies block COVID-19 virus binding to its receptor ACE2 doi: https://doi.org/10.1101/2020.05.01.20077743

48. Barde I, Verp S, Offner S, Trono D. Lentiviral Vector Mediated Transgenesis. Curr Protoc Mouse Biol. 2011 Mar 1;1(1):169–84. doi: 10.1002/9780470942390.mo100169.

49. Hoogenboom HR, Griffiths AD, Johnson KS, Chiswell DJ, Hudson P, Winter G. Multi-subunit proteins on the surface of filamentous phage: methodologies for displaying antibody (Fab) heavy and light chains. Nucleic Acids Res. 1991 Aug 11;19(15):4133–7.

50. Kingsbury GA, Junghans RP. Screening of phage display immunoglobulin libraries by anti-M13 ELISA and whole phage PCR. Nucleic Acids Res. 1995 Jul 11;23(13):2563–4.

51. Yuan QA, Simmons HH, Robinson MK, Russeva M, Marasco WA, Adams GP. Development of engineered antibodies specific for the Müllerian inhibiting substance type II receptor: a promising candidate for targeted therapy of ovarian cancer. Mol Cancer Ther. 2006 Aug;5(8):2096–105.

52. Hong P, Koza S, Bouvier ES. Size-Exclusion Chromatography for the Analysis of Protein Biotherapeutics and their Aggregates. J Liq Chromatogr Relat Technol. 2012;35(20):2923–2950. doi: 10.1080/10826076.2012.743724

